# Integrative cell bin segmentation on spatial transcriptomics by Voronoi

**DOI:** 10.1101/2024.11.07.622434

**Authors:** Ming Lin

## Abstract

Spatial transcriptomics is undergoing rapid advancements and iterations. It is a beneficial tool to significantly enhance our understanding of tissue organization and relationships between cells. Recent technological advancements have achieved subcellular resolution, providing much denser spot placement for downstream analysis. A key challenge for this following analysis is accurate cell segmentation and the assignment of spots to individual cells.

The primary objective of this study was to evaluate the effectiveness of a new cell segmentation approach based on subcellular level spatial transcriptomic data by confirming nuclei positions and using Voronoi diagrams, compared to direct clustering with cellbin data. Our findings demonstrate that the Voronoi method not only outperforms traditional methods in providing clearer boundaries and better separation of cell types but also excels in preserving the most transcripts, addressing the issue of low capture efficiency. This integrative methodology presents a substantial advancement in spatial transcriptomics, offering improved cell type classification and spatial pattern recognition. All codes in the paper are available at the GitHub repository: https://github.com/Charlottttttte/Cell_Segmentation_Voronoi

## Introduction

Spatial transcriptomics can significantly enhance our comprehension of tissue arrangement and intercellular communications(1,2), preserving spatial information lost in traditional single-cell RNA sequencing (scRNA-seq)(3). This information is significant for analyzing cancer(4), finding relationships between different genes(5), and discovering drugs(6). In current biomedical research, Spatial transcriptomic technologies can be generally divided into two categories, imaging-based technologies (IST) and sequencing-based technologies(SST)(7). Imaging-based methods, such as MERFISH(8), seqFISH+(9), can obtain high-resolution gene expression information but are limited in their ability to detect all gene types. In contrast, sequencing-based methods which benefit by their efficiency and high throughput, originally faced the challenge of low resolution. These initial SST methods such as Visium(10) only have 55 μm (center-to-center) capture areas, capturing gene expression from multiple cells within each spatial area, making it difficult to discern single-cell level details(11).

Recent advancements have addressed some of these limitations. For instance, the development of Slide-seqV2^11^ has improved the spatial resolution of SST to 2 μm, enabling near single-cell resolution by capturing transcripts with higher efficiency and spatial precision. The advent of the Stereo-seq (Spatial Enhanced Resolution Omics Sequencing) method from BGI Spatial has further enhance the capture density to 0.5 μm(center-to-center). Stereo-seq integrates high-resolution spatial barcoding with next-generation sequencing, enabling the capture of gene expression at subcellular resolution(12), demonstrating the highest capturing ability among the current SST method(13). This improvement allows for precise mapping and analysis of gene expression within individual cells, significantly improving our ability to study the intricate spatial dynamics of tissues and facilitate a deeper understanding of cellular functions.(14)

Despite these advancements, critical analysis reveals that current methods still have room for improvement(13). IST methods, while offering high resolution, require complex and time-consuming procedures. SST methods, despite enhancements in resolution, still struggle with the comprehensive capture of spatial context at a single-cell level(15). Therefore, there is a continuous need for innovative approaches that combine the strengths of both IST and SST method to achieve more precise and comprehensive spatial transcriptomic analyses(16).

When we obtain the gene expression information by the above methods, accurate cell segmentation is essential before we ultimately cluster the cells together and do downstream analyses(17). The challenge of attaining precise, automated cell segmentation primarily stems from variations in cell morphology, dimensions, and distribution within different tissue types(18).

Conventional image-based segmentation method in sequencing-based technologies are constrained and fail to harness the information provided by spatial transcriptomics profiling fully(18). Some original method use watershed algorithm(19) to find the cell boundaries, other recent method design deep learning-based cell segmentation algorithm to handle complex tissue images, including TissueNet(20), GeneSegNet(21), Cellpose(22), and SCS(23). TissueNet designs its primary function to accurately identify and segment cells in high-dimensional tissue images, leveraging a neural network trained on a diverse set of annotated images to generalize well across different tissue types and imaging modalities.

For high-resolution spatial transcriptomics, SCS was designed by integrating sequencing and imaging data, utilizing transformer neural network to adaptively learn the position of each spot relative to its cell center, finally enhancing cell segmentation and achieving greater accuracy compared to current methods. However, those methods still face some drawbacks due to the requirement of supervision(24), low capture efficiency and lengthy code runtimes(25).

In this study, we introduce the Voronoi method(26), which is designed on nuclei-based data and contains larger gene expression data. Built on the BGI method, our Voronoi method will optimize their cell segmentation method by utilizing Voronoi segmentation(27) combined with nuclei-based spatial data, providing more accurate cell type clustering and larger dataset that preserves most transcripts for downstream analysis.

This method uses spatial transcriptomics data, especially cellbin.gef and tissue.gef data. First, we utilized cellbin data to determine the exact location of nuclei and define them as the center of each cell region, then the Voronoi diagram was employed to delineate cell boundaries, assigning each nucleus to a unique region index. We assume each region represents a cell, gene information from tissue data was then mapped onto these segmented regions, obtaining a larger cellbin data consisting of fourfold numbers of gene expression. Clustering and further downstream analysis were performed by using the Stereopy toolkit(28) and Mapmycells(29) to prove the dominant advantage of Voronoi method.

## Results

To validate the effectiveness of the Voronoi segmentation method, we first applied it to a small section of mouse brain tissue, consisting of 72 cells. This initial test demonstrated the feasibility of our approach on a manageable scale. Encouraged by these results, we expanded the analysis to a 3000×3000 µm section of the mouse brain to assess the impact of integrating tissue data on a larger and more complex dataset. The Voronoi method continued to demonstrate robust performance, effectively delineating cell regions and integrating genetic information with greater precision than the original cellbin dataset.

### The generation of Voronoi

The generation of the Voronoi diagram was a pivotal step in our analysis. By using the spatial coordinates of nuclei from the cellbin data, we defined the centers of each cell region, which the Voronoi method then used to delineate boundaries (**Fig 1a**). This approach ensured that each nucleus was assigned a unique region, thereby accurately representing individual cells. By using the Voronoi Segmentation method, we cut the mouse brain section into certain regions that own the same number as the numbers of nuclei(cells), as shown in **fig1b**. Cartesian coordinates of vertices and boundaries can be obtained so that each region has its individual range presented in cartesian coordinates and is assigned a unique region index (**Fig 1c**).

**Fig. 1.**
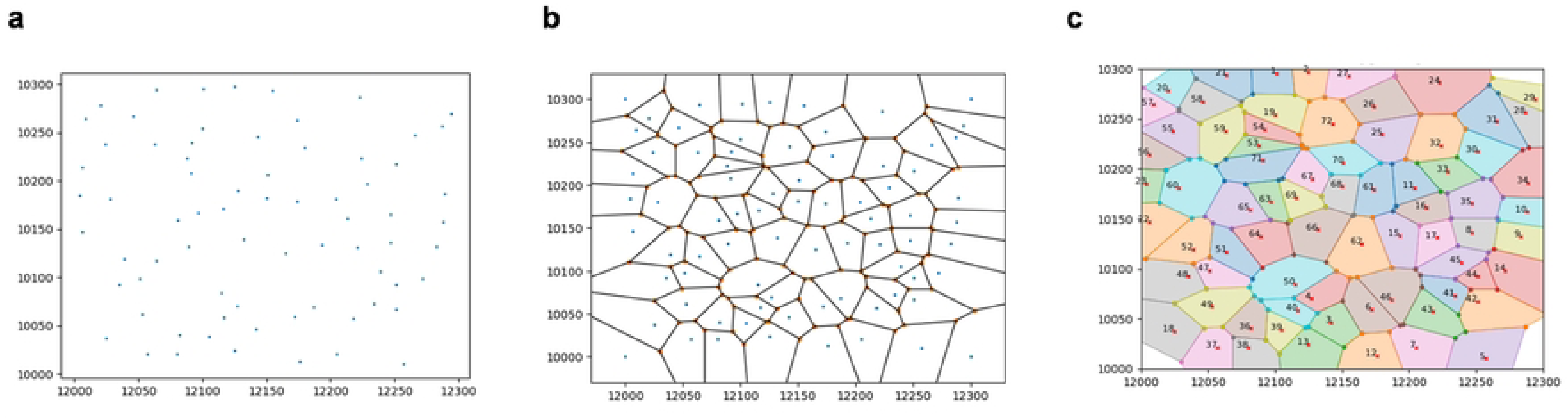
Visualization of Cell Locations and Voronoi Segmentation method. (a) The cell locations obtained from the cell.gef file is shown in this raster plot, illustrating a specific region for 72 nuclei. The x-axis ranges from 12000 to 12300, and the y-axis ranges from 10000 to 10300. Each blue dot represents an individual cell, which serves as the center point for the Voronoi segmentation method. (b) This diagram demonstrates the Voronoi segmentation applied to the cell locations. Each solid black line represents the boundary of a cell region, generated such that any point within a region is closer to its corresponding cell (center point) than to any other cell. (c) This visualization assigns unique region indices to each Voronoi cell. Different colors represent distinct regions, and each region is labeled with a specific index.

### Integration of tissue data

Recognizing that cellbin data primarily includes information about the locations of cell nuclei as detected during staining, it represents only a small fraction of the spatial landscape within the tissue slice. This limitation restricts the comprehensive analysis of gene expression across the entire tissue. In contrast, tissue data encompasses the entire tissue slice, providing a broader and more detailed spatial transcriptomic profile. To overcome the spatial limitations of cellbin data, we integrated tissue data with the cellbin nuclei locations. By mapping the denser tissue data onto the regions defined by the cellbin nuclei, we effectively quadrupling the spatial resolution and gene expression information within each region, resulting in a more complete dataset that captured gene expression across the entire tissue slice.

### Use Stereopy to process and analyze the dataset

After integrating tissue data into each region using the Voronoi method, we created a new, enriched dataset—referred to as the “Voronoi dataset.” This dataset offers a more comprehensive view of gene expression across the entire tissue section. To understand the extent of the improvements introduced by the Voronoi method, we compared this new dataset with the original cellbin dataset, which we’ll refer to as the “Original dataset.”

Both datasets maintain the same structural format: an information matrix where rows correspond to spatial positions and columns represent different gene types. To evaluate the impact of the Voronoi method, we used the BGI toolkit to process and analyze, aiming to objectively assess whether the Voronoi method enhances the clustering quality and spatial resolution of the resulting analysis.

### Comparison between the Voronoi dataset and Original dataset

Our analysis began with a 3000×3000 μm section of the mouse brain, carefully selected to test the method’s effectiveness in a complex tissue environment. As illustrated in **Fig 2**, we compared the spatial distribution of clusters identified using the Leiden clustering method after PCA.

**Fig. 2.**
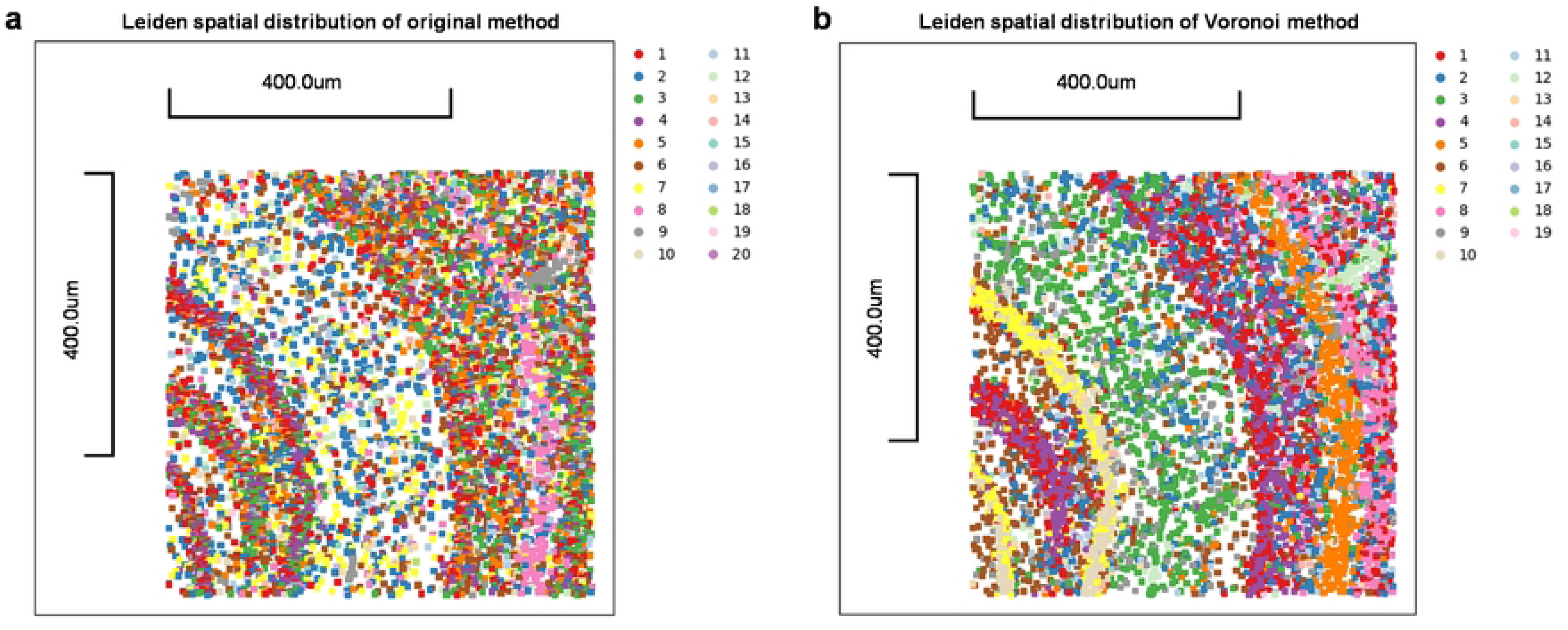
Comparison of Leiden clustering results using different cell segmentation methods in spatial transcriptomics. (a) Leiden spatial distribution of the original method. (b) Leiden spatial distribution of the Voronoi method. Colors represent different clusters as identified by the Leiden algorithm. The scale bar represents 400.0 μm in (a) and (b).

**Fig 2a** displays the clustering results from the Original dataset, with each color corresponding to a specific cluster as indicated in the legend. The clusters, while distinguishable, show some overlap and blurred boundaries, indicating challenges in accurately defining cell types and spatial regions. This limitation is particularly evident in the central regions of the tissue, where different cell types are densely packed.In contrast, **Fig 2b** shows the clustering results after applying the Voronoi method. These clusters are more refined and distinct, with clearer separation between different cell types.

**Figs 3a** and **3b** further highlight the method’s impact on specific clusters. In the Original dataset, clusters appeared dispersed with indistinct boundaries, complicating conclusions about spatial organization. However, the Voronoi method significantly improved the clarity and concentration of clusters, aligning well with known anatomical structures in the mouse brain, underscoring its accuracy in cell classification.

**Fig. 3.**
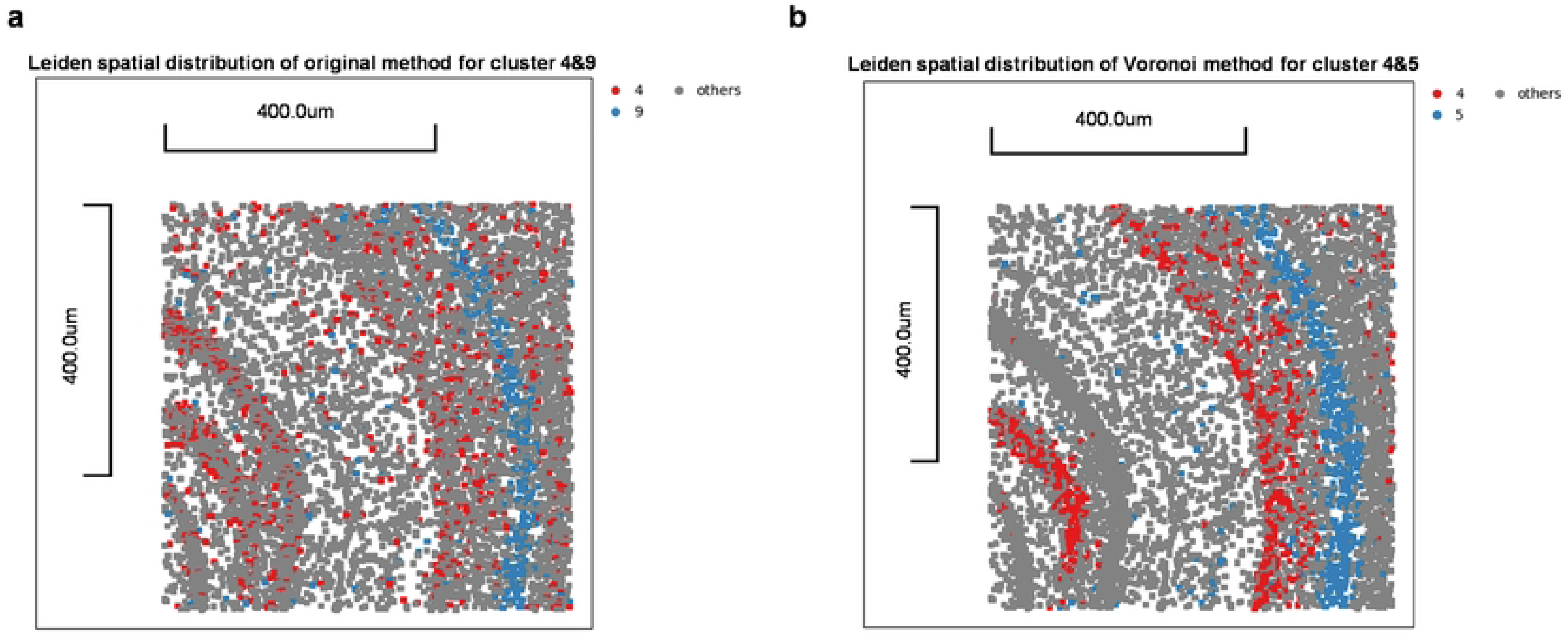
Comparison of Leiden clustering for specific clusters. (a) Leiden spatial clus-tering of original method for specific cluster 4 & 9, others are shown in grey. (b) Leiden spatial clustering of Voronoi method for specific cluster 4 & 5, others are shown in grey. Red and blue dots represent two clusters identified by the Leiden algorithm in the similar layer. The scale bar represents 400.0 μm both in (a) and (b).

### UMAP Embeddings and Clustering Quality

Moving to a broader analysis, **Fig 4** compares UMAP embeddings from both methods. The Voronoi method consistently outperforms the Original method, producing more distinct and less overlapping clusters. This result indicates a more accurate identification of cellular identities, which is critical for understanding the complex spatial dynamics within tissues.

**Fig 4:**
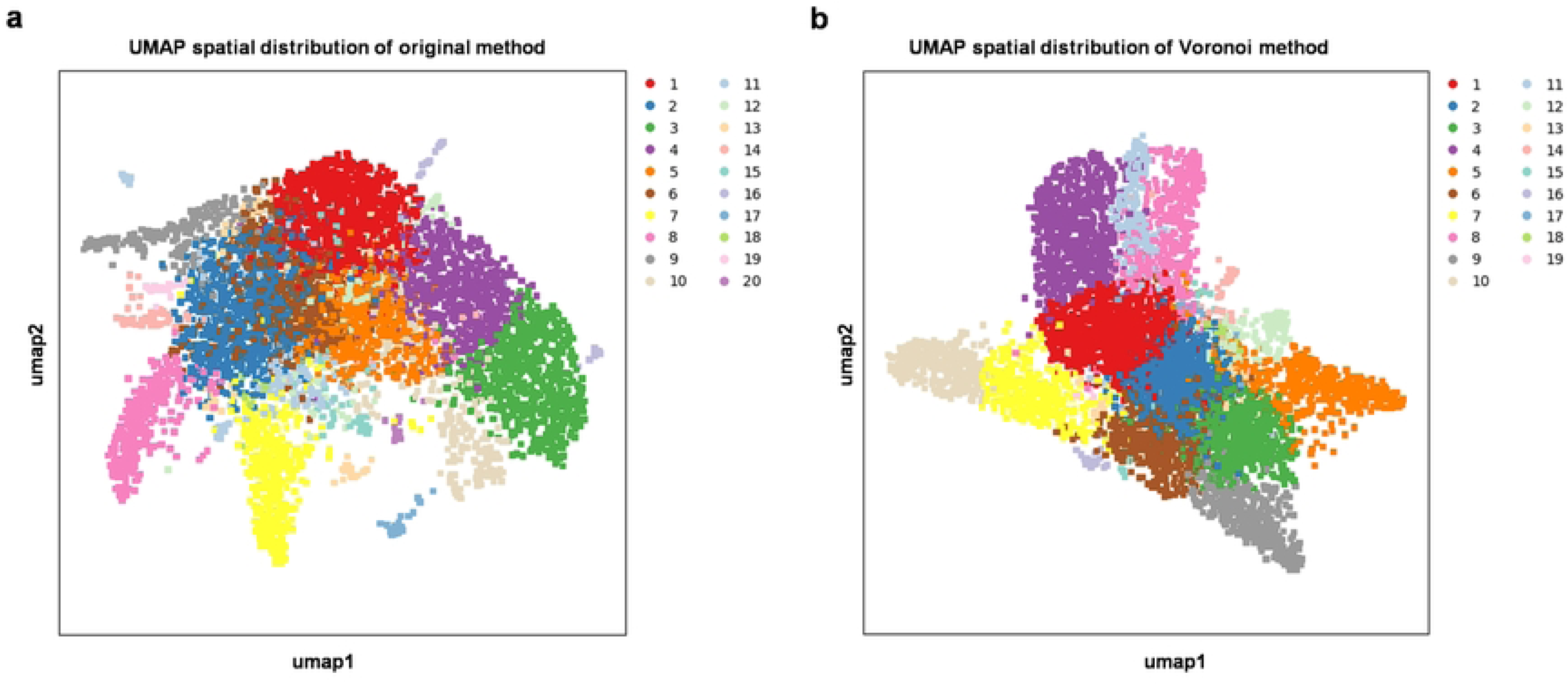
Comparison of Umap spatial distribution using different method. (a) UMAP spatial distribution of the original method. (b) UMAP spatial distribution of the Voronoi method. Both panels display the UMAP embedding of the clustering results obtained using the original and Voronoi method. Each point represents a cell, and the colors correspond to different clusters as identified by the Leiden algorithm.

### Quantitative Analysis of Clustering Quality

To quantify the improvements, we calculated the Silhouette Score and Calinski-Harabasz Index for both methods, where higher values in both indices denote more distinct and well-defined clusters. The Voronoi method achieved a significantly higher Silhouette Score (0.166) and Calinski-Harabasz Index (2474.823) compared to the Original method (0.113 and 1256.078, respectively). These enhancements confirm that the Voronoi method produces more coherent and distinct clusters, reflecting better spatial separation and cluster compactness.

Moreover, the Voronoi method captured a higher total count of genes and identified more non-zero gene counts within each cell, as shown in **Table 1**. These comparisons suggest that the Voronoi method not only improves detection of gene expression diversity but also enhances spatial resolution and clustering result, offering a more detailed and precise view of the tissue’s molecular landscape.

**Table 1:**
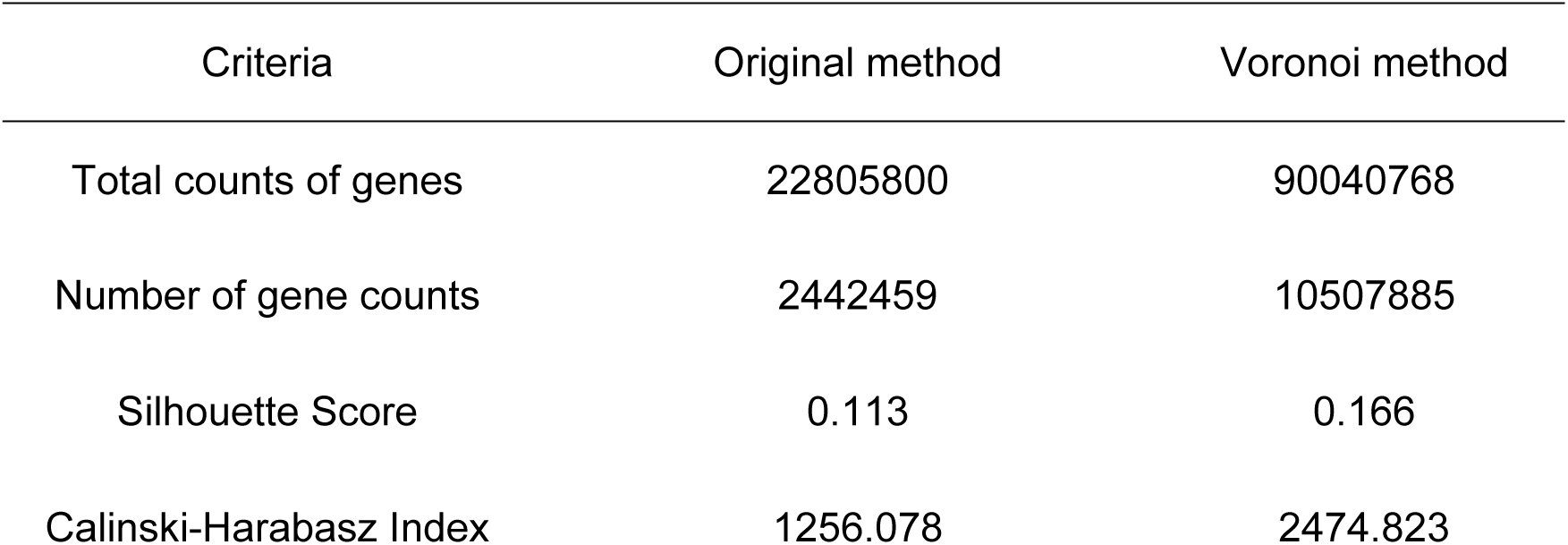
Comparison of dataset and clustering results between original and Voronoi methods.

| Criteria | Original method | Voronoi method |
| --- | --- | --- |
| Total counts of genes | 22805800 | 90040768 |
| Number of gene counts | 2442459 | 10507885 |
| Silhouette Score | 0.113 | 0.166 |
| Calinski-Harabasz Index | 1256.078 | 2474.823 |

In conclusion, the Voronoi segmentation method demonstrates substantial improvements in cell type classification and spatial analysis compared to traditional approaches. It provides a more accurate and clearer representation of spatial layers in transcriptomic data, aligning better with known anatomical structures. This method not only aligns better with the known anatomical structures of the mouse brain as supported by Allen Brain Atlas(30), but also holds promise for future applications in other tissues, paving the way for deeper insights into cellular function and tissue organization.

## Mapmycells method Analysis

### Comparison of Clustering Quality

To further evaluate the effectiveness of the Voronoi method in analyzing spatial transcriptomics data, we employed the MapMyCells(29) tool. This tool is designed to map scRNA-seq and spatial transcriptomics data to cell types with bootstrapping probability, enabling scientists to compare their transcriptomics and spatial data against Allen Institute’s datasets. We analyzed both the original and Voronoi-processed datasets to evaluate how well each method captured the spatial distribution of cell types across a full brain tissue section.

### Overall Spatial Distribution Analysis

Initially, a smaller region (3000*3000 μm) was analyzed, but to better learn the distribution of cell types and capture more contextual information across the entire mouse brain, the analysis was expanded to a full brain tissue section. The following **fig 5** shows scatter plots representing entire mouse brain data processed by MapMyCells and categorized by cell type. Each plot visualizes the distribution of cell class within the brain slice, with distinct colors indicating various cell types as indicated by the class number on the right. This figure allows for a comparison of the clustering quality between the original method and the Voronoi method.

**Fig. 5.**
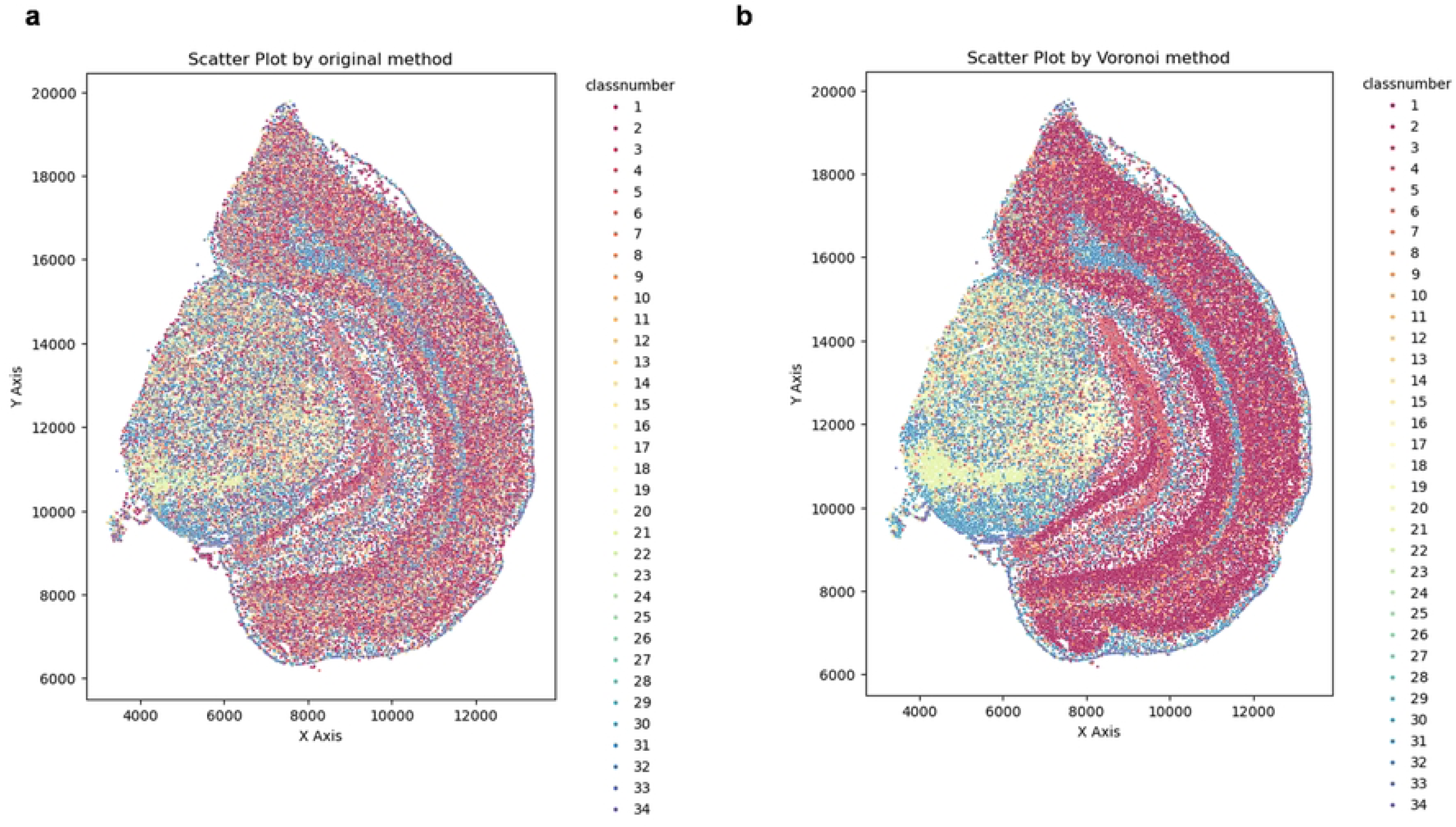
Comparison of mouse brain slice scatter plots after MapMyCells process. (a) Scatter plot by original method. Each color represents a class of cell type, ranging from class 1 to class 34. (b) Scatter plot by Voronoi method. Each color represents a class cell type, ranging from class 1 to class 34. The X and Y axes represent spatial coordinates within the brain slice.

**Fig 5a** shows the results from the Original dataset. Here, the scatter plot reveals some overlap between clusters, particularly in the central region, where different cell types are mixed. This overlap suggests that the Original method may struggle to clearly separate different cell types, leading to potential inaccuracies in cell type identification and spatial distribution analysis. In contrast, **Fig 5b** shows the Voronoi method, with less overlap and more distinct clusters, indicating improved clustering performance. The Voronoi method’s enhanced separation between clusters allows for a clearer and more accurate spatial representation, which is critical for understanding cell interactions and functions within the tissue.

### Detailed Analysis of Layer 2

Mouse brain’s cerebral cortex is divided into six layers, each serving distinct functions and composed of various cell subtypes based on their spatial positions(31). To further evaluate clustering quality, we focused on Layer 2, which has a unique cellular composition. MapMyCells classified cells into specific subtypes, and the spatial distribution of these subtypes was compared against known anatomical references to assess dataset quality.

**Fig 6** consists of two scatter plots illustrating the distribution of unique cell subtypes that belong to layer 2 with bootstrapping probability. The cell subtypes represented in the plots can be regarded as marker subtypes of layer 2(32).

**Fig. 6:**
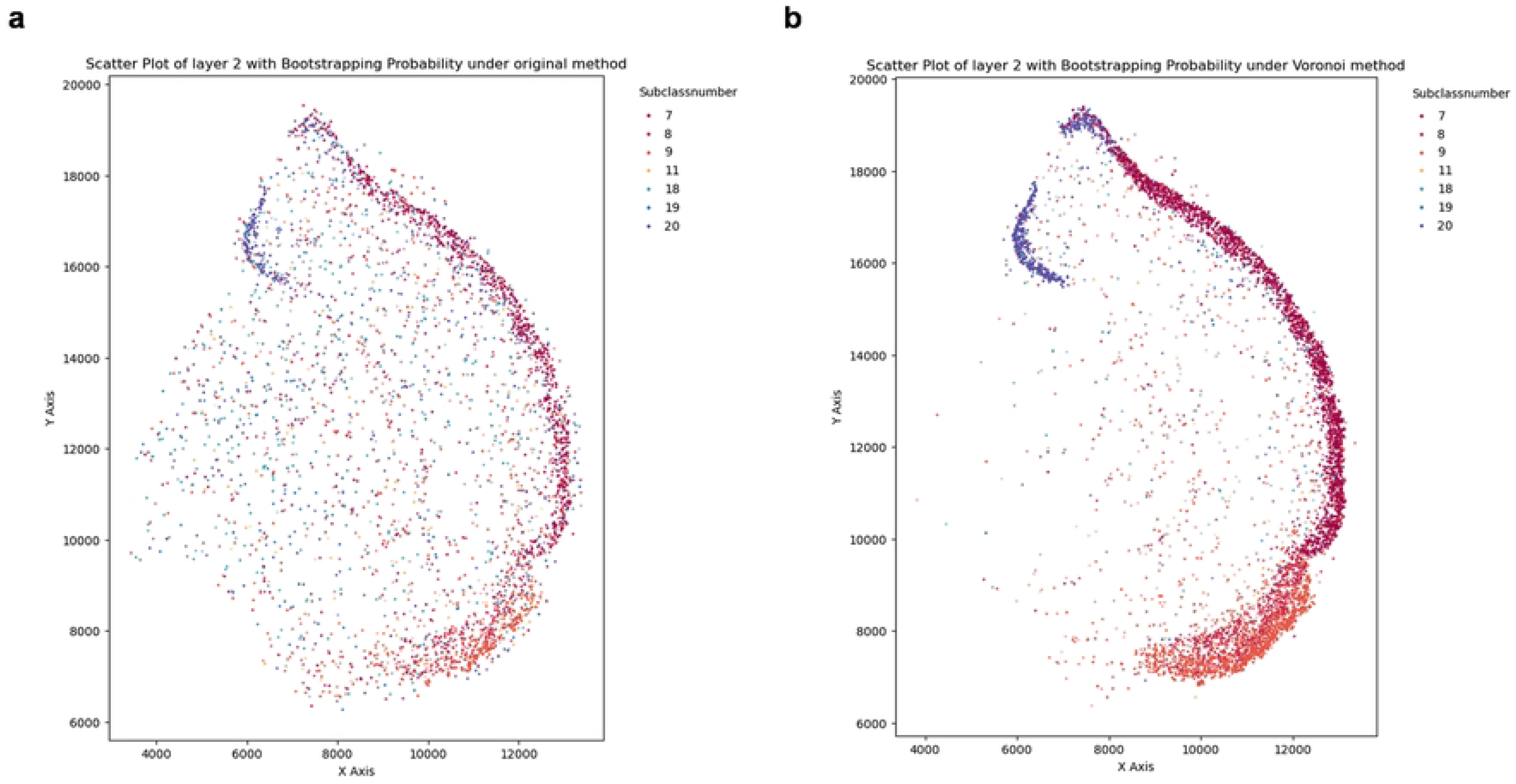
Comparison of spatial distribution of layer 2 in entire slice by Mapmycells. (a) Scatter Plot of layer 2 by original Method. (b) Scatter Plot of layer 2 by Voronoi Method. Each point represents a cell, with colors corresponding to different subtypes. The transparency of each point indicates confidence in cell subtype classification, with darker points signifying higher probabilities.

**Fig 6** illustrates the distribution of key marker subtypes for Layer 2 detected in both datasets. The Voronoi method (**Fig 6b**) displays a denser and more defined boundary for Layer 2 compared to the Original method (**Fig 6a**), which shows lighter and less distinct boundaries with significant noise. The improved clarity and boundary definition in the Voronoi dataset suggest higher confidence in subtype classification, reflected in the darker points indicating higher bootstrapping probabilities. The reduction of noise further supports the Voronoi method’s superior simulation of the ground truth distribution, reinforcing its effectiveness in accurately mapping cellular subtypes.

### Quantified Analysis of Layer 5 and Gaussian Fitting Comparison

To further assess the sharpness performance of the Voronoi method in comparison to the Original method, we conducted a detailed analysis on Layer 5 of the mouse brain. Instead of using the standard x-axis or y-axis for analysis, we selected a custom axis because direct analysis along either axis fails to capture the true spatial distribution of cells.

Using this custom axis, we measured the distribution of cells and evaluated the fit of the data to the custom line to quantify how well each method captured the non-linear spatial dynamics of cell distributions. **Figs 7a** and **7b** present scatter plots of the cells in Layer 5 using the Original and Voronoi methods, respectively, along with the custom line fitted to the data. This custom axis, shown in red, is represented by the equation *y* = ―0.85x + 22400, and serves as a reference for analyzing the spatial organization of the cells relative to this axis.

**Fig. 7.**
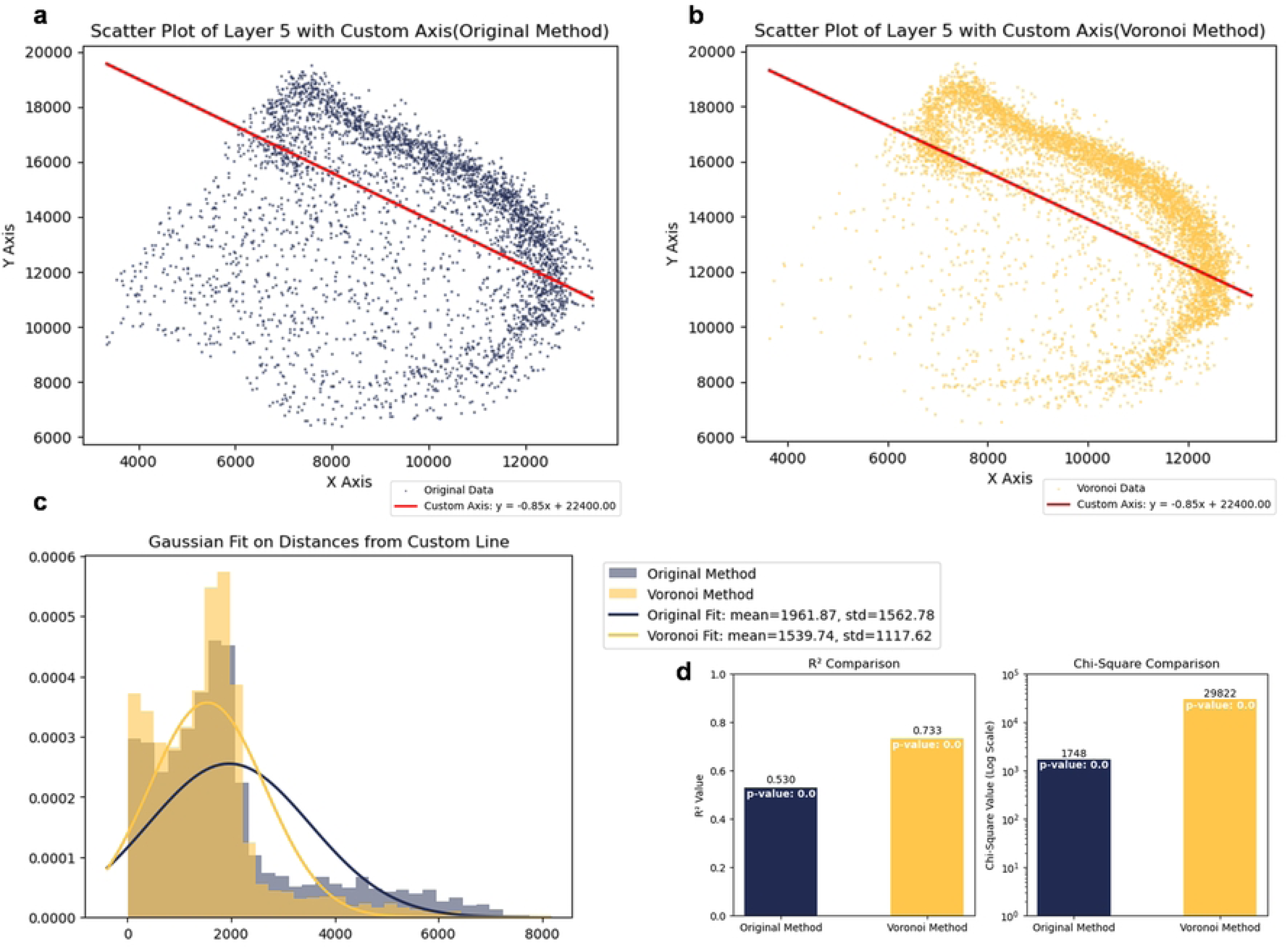
Quantified Analysis of Layer 5. (a) Scatter plot of Original Method with custom axis: The distribution of cells in Layer 5 using the Original method, plotted along a custom axis (red line) represented by the equation y=-0.85x+22400. (b) Scatter plot of Voronoi Method with custom axis: The cell distribution in Layer 5 using the Voronoi method, with the same custom axis. (c) Gaussian fit of distances from the custom axis: A frequency plot showing the distribution of distances of cells from the custom axis, with Gaussian fitting curves for both methods. (d) Bar plot comparing R² and Chi-square values: Comparison of the R² values and Chi-square statistics for the Gaussian fits of the Original and Voronoi methods.

To quantify the differences between the two methods, we measured the distances of each cell from the custom line and performed a Gaussian fit on the distribution of these distances, as shown in **Fig 7c**. The frequency plot illustrates the distribution of distances for both methods, with the Original method represented by the blue curve and the Voronoi method by the orange curve.

The Gaussian fitting results reveal that the Voronoi method produces a narrower distribution, with a lower mean distance (1539.74) and a smaller standard deviation (1117.62), compared to the Original method (mean = 1961.87, std = 1562.78). This indicates that the Voronoi method aligns the cells more closely with the custom axis, further supporting its superior ability to capture the true spatial dynamics of the tissue.

Finally, **Fig 7d** presents a comparison of two key statistical metrics: the R² value and the Chi-square value, both of which assess the goodness-of-fit for the Gaussian distributions. The bar plots on the left side of the figure show that the Voronoi method achieves a significantly higher R² value (0.733) compared to the Original method (0.530), indicating a better fit to the custom line. On the right, the Chi-square test results also favor the Voronoi method, with a substantially lower Chi-square value (1748) compared to the Original method (29822), further confirming the improved performance of the Voronoi method.

## Methods

### Data Source

The data used in this study were obtained from Stereopy, a Python package developed by BGI Spatial for spatial transcriptomics analysis. The main dataset, the MouseBrain Demo, includes two GEF files: SS200000135TL_D1.cellbin.gef and SS200000135TL_D1.tissue.gef. GEF files provide spatial coordinates, gene expression data, and metadata crucial for understanding spatial gene expression. Cellbin data represent gene expression at the cell nucleus level, while tissue data include expression data for spots within the mouse brain tissue. The dataset can be downloaded from the following URL: Steropy Tutorial Data.

### Voronoi Segmentation Method

The Voronoi diagram was constructed using nuclei coordinates as seed points, utilizing Euclidean distance to partition the tissue into distinct cell regions. Each Voronoi cell represented a unique region assigned to its corresponding nucleus, delineating clear cell boundaries based on proximity criteria.

### BGI Data Processing Method

The **BGI process** consists of Preprocessing, Embedding, and Clustering:

#### Preprocessing

Preprocessing is crucial for ensuring data quality and preparing it for subsequent analysis. The preprocessing steps are shown as follows:

**1. Data filtering**: To ensure accuracy and reliability of the analysis, we make the quality control: remove all missing values and outliers from the dataset, as well as cells having too many mitochondrial genes expressed, cells without enough genes expressed, and cells exceed the count range. Here we delete the cells whose number of genes that have non-zero counts are less than 3.
**2. Normalization**(33): Scaling the data to ensure that all features contribute equally to the analysis. This involves adjusting the values measured on different scales to a common scale. Methods for normalization are normalize_total(34) and log1p(35).
**3. Filtering**: Highly variable genes are then selected based on predefined criteria to focus the analysis on the most pertinent features. The steropy package provides preloaded tools to handle and preprocess such spatial datasets effectively.

#### Embedding

Embedding refers to transforming high-dimensional data into a lower-dimensional space to facilitate visualization and analysis:

**1. Dimensionality Reduction**: Techniques such as PCA are applied to reduce the dimensionality of the data while preserving its intrinsic structure. Only highly variable genes are taken into consideration in this step. After that, we calculate the neighborhood graph(36) of cells and use UMAP method(37) with the help of PCA representation of the expression matrix.
**2. Visualization**: The lower-dimensional embeddings are visualized to observe patterns and relationships within the data. UMAP is used to help in understanding the data’s underlying structure and distribution.

#### Clustering

Clustering is the process of grouping similar data points together based on their features:

**1. Algorithm Selection**: Leiden algorithm, which has been proved to fit spatial transcriptomics better than Louvain, is selected(38).
**2. Cluster Assignment**: Each data point was assigned to a cluster, and the resulting clusters were analyzed to understand their characteristics. Clusters can be visualized in scatter plots and clustering effects can be evaluated by UMAP.

### Clustering Evaluation Method

#### Silhouette Score(**39**)

The Silhouette Score measures the cohesion and separation of clusters. It quantifies how similar each data point is to its own cluster compared to other clusters. The score ranges from -1 to 1, where a higher value indicates better-defined and more distinct clusters.

The Silhouette Score for point *i* is defined as: 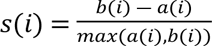

#### Calinski-Harabasz Index(**40**)

The Calinski-Harabasz Index, also known as Variance Ratio Criterion, evaluates the ratio of between-cluster variance to within-cluster variance. Higher values indicate better-defined clusters.

Set *k* as the number of clusters, *n* as the total number of data points, *S*_*B*_ be the between-cluster dispersion matrix, and *S_w_*be the within-cluster dispersion matrix.

The index is computed as: 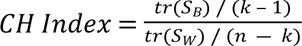

Here, _tr_ (*S*_*B*_) is the trace of the between-cluster dispersion matrix and _tr_ (*S*_w_) is the trace of the within-cluster dispersion matrix.

### Single-Cell Data Analysis with MapMyCells

Single-cell transcriptomic data were analyzed using MapMyCell(29), a comprehensive software suite designed for the integration, visualization, and interpretation of single-cell RNA sequencing (scRNA-seq) data. The software supports the integration of multiple datasets and allows for the comparison of cell populations across different conditions or treatments. Key features include customizable visualization options, such as t-SNE and UMAP plots, which facilitate the identification of distinct cell types and states.

## Conclusion

The comparison between the clustering capability from the original method and the Voronoi method proves that the Voronoi method significantly improves cell type clustering quality. The Voronoi method achieves better separation and clearer boundaries between cell types, as well as more compact and coherent clusters, and sharper distributions. These improvements are both visually and quantitatively evident and support the use of the Voronoi method for more accurate and reliable cell type clustering in spatial transcriptomics data.

